# Look4TRs: A *de-novo* tool for detecting simple tandem repeats using self-supervised hidden Markov models

**DOI:** 10.1101/449801

**Authors:** Alfredo Velasco, Benjamin T. James, Vincent D. Wells, Hani Z. Girgis

## Abstract

Simple tandem repeats, microsatellites in particular, have regulatory functions, links to several diseases, and applications in biotechnology. Sequences of thousands of species will be available soon. There is immediate need for an accurate tool for detecting microsatellites in the new genomes. The current available tools have limitations. As a remedy, we proposed Look4TRs, which is the first application of self-supervised hidden Markov models to discovering microsatellites. It adapts itself to the input genomes, balancing high sensitivity and low false positive rate. It auto-calibrates itself, freeing the user from adjusting the parameters manually, leading to consistent results across different studies. We evaluated Look4TRs on eight genomes. Based on F-measure, which combines sensitivity and false positive rate, Look4TRs outperformed TRF and MISA — the most widely-used tools — by 106% and 82%. Look4TRs outperformed the second best tool, MsDetector or Tantan, by 11%. Look4TRs represents technical advances in the annotation of microsatellites.

## Introduction

Genomes contain an abundance of repeated sequences known as simple tandem repeats, each of which consists of a motif repeated in tandem. Due to mutations and replication errors, motif copies may vary remarkably. Simple tandem repeats are classified according to the length of the motif into microsatellites (MS), minisatellites, and satellites^1^. Simple tandem repeats — MS in particular — have important functions including gene regulation and recombination^2–6^. Further, MS have been linked to many diseases such as colon cancer, Fragile X syndrome, myotonic dystrophy, Kennedy’s disease, Huntington’s disease, diabetes, and epilepsy^1, 7–13^. Furthermore, MS are utilized in paternity tests and forensic medicine^14^. Moreover, in the bioinformatics field, excluding MS while aligning sequences has been reported to improve the performance of alignment algorithms^15^.

Recent advances in sequencing technology have led to multiple, large-scale sequencing projects such as The Cancer Genome Atlas (https://cancergenome.nih.gov), the 100,000 Genomes Project (https://www.genomicsengland.co.uk), the 1000 Genomes Project (http://www.internationalgenome.org), the Genome 10K Project for vertebrates^16^, the i5K Project for insect genomes (http://i5k.github.io), the 10KP Project for plant genomes (https://db.cngb.org/10kp/), and the Earth Biogenome Project (https://www.earthbiogenome.org).

Advances in sequencing technology has outpaced advances in annotating the new genomes, including the annotation of MS. Multiple computational tools have been developed for detecting MS. RepeatMasker (http://www.repeatmasker.org) is a widely used for locating all types of repeats — tandem and interspersed. This tool searches for instances of a manually-curated consensus sequence in a genome. RepeatMasker cannot identify instances of unknown MS. Alternatively, de-novo tools can identify MS regardless if their motif is known or not^13, 15, 17–19^.

However, the currently available *de-novo* tools tends to fall into two categories. Tools that are very sensitive to MS but produce high false positive rates belong to the first category, whereas tools that are not sensitive but produce low false positive rate belong to the second category. We believe that these two performance extremes are due to disregarding the information available in an input genome. Generally, tools are tested on certain genomes with certain properties. If a tool is given a new genome with properties unlike the tested genomes and this tools were incapable of evaluating the content of the new genome, it would most likely fail to perform well. For example, consider the *Plasmodium falciparum* genome (the parasite causing malaria in humans), where 80% of its nucleotide composition is composed of AT. If a tool did not consider this biased nucleotide composition, its false positive rate would be very high. RepSeek is the first tool for detecting interspersed repeats while accounting for the nucleotide composition of the input sequences^20^. Inspired by RepSeek, Red takes into account the k-mer composition of a genome while annotating repeats^21^. We are not aware of any MS-detection tool that accounts for sequence composition. Yes, MS-detecting tools — except MsDetector — provide adjustable parameters that can be used to adapt a tool to the input genome (MsDetector can be retrained on an input genome). However, adjusting these parameters is impractical given the thousands of genomes being currently sequenced.

Therefore, there is an immediate need for a auto-calibrating, adaptive tool for annotating MS in hundreds of thousands of new genomes. To this end, we propose Look4TRs, which is a novel, sophisticated, computational tool that remedies the above mentioned limitations. Through Look4TRs, we make three main contributions to the methodology of MS discovery. First, Look4TRs takes into account the nucleotide composition of the input system. Second, Look4TRs is the first self-supervised system for annotating MS. Third, Look4TRs is the first auto-calibrating system for locating MS. Supervised learning algorithms require the availability of labeled examples, whereas self-supervised learning algorithms can generate its own training labeled examples. Look4TRs utilizes self-supervised hidden Markov models (HMMs) and self-supervised general linear models. Tantan^15^, which is a tool for detecting MS, utilizes an HMM. The HMM parameters are set manually. MsDetector^22^, which is another tool for detecting MS, utilizes an HMM too. The HMM parameters are determined using known MS located by RepeatMasker, i.e. these parameters are set automatically; this process has to be repeated on each genome for optimal performance. Look4TRs generates a random chromosome based on a real chromosome of the input genome. Then it inserts semi-synthetic MS in the random chromosome. Finally, the HMM is trained and calibrated using these semi-synthetic MS. Three parameters affect the training of Look4TRs’s HMM. Look4TRs automatically trains multiple HMMs using all valid combinations of these three parameters. The HMM resulting in the best balance between high sensitivity and low false positive rate is selected automatically. These features enable Look4TRs to adapt to the input genome automatically. Therefore, Look4TRs represents a true progress in the methods for MS discovery.

## Results

### Evaluation measures

Look4TRs, Tantan^15^, TRF^17^, MISA^18, 19^, and MsDetector^22^ are evaluated using the following eight criteria: (i) sensitivity, (ii) False Positive Rate (FPR), (iii) precision, (iv) F-measure, (v) novel microsatellites (MS) content, (vi) percentage coverage, (vii) time requirement, and (viii) memory requirement. Let’s define the true positives (TP) as the number of nucleotides that comprise MS found by RepeatMasker and by a tool and the false negatives (FN) as the number of nucleotides of MS found by RepeatMasker but missed by a tool. The false positives (FP) is the number of nucleotides found by a tool in a random synthetic genome that has the same length and nucleotide composition as the input genome; simple repeats found by RepeatMasker in the random synthetic chromosome are removed. Equations 1 and 2 define the sensitivity and the precision in terms of TP, FP, and FN.

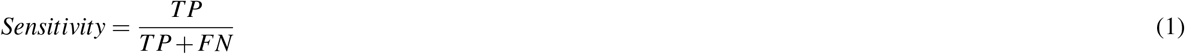

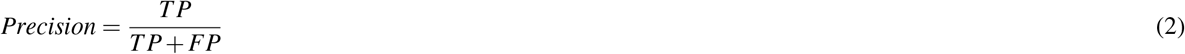

The FPR is the total length of false positives in 1 mega base pair (mbp) and defined by Equation 3.

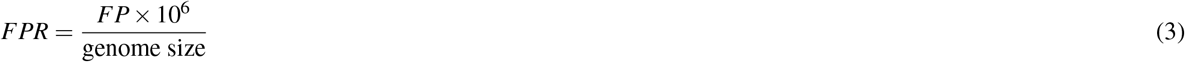

A combination of precision and sensitivity defines the F-measure (Equation 4), which reflects how well a tool balances between sensitivity and precision.

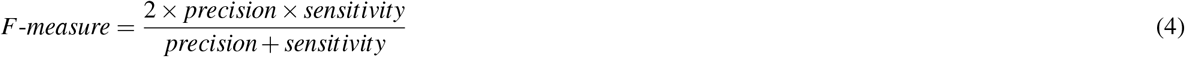

Repbase is a database of manually annotated repeats^23, 24^. RepeatMasker is an alignment-based scanning tool, which utilizes Repbase. If a repeat sequence is not in Repbase, RepeatMasker will not be able to find it. The novel MS content is the number of neocleotides comprising potential MS that are not found by RepeatMasker. Another evaluation measure is the percentage coverage, which is defined as the percentage of MS located by a tool in the input genome. We measured the time and the memory requirements on a laptop with a 2.8 GHz Intel Core i7 processor and 16 GB RAM.

**Table 1.** Comparison among five MS-detection tools on eight genomes. Sensitivity measures the percentages of nucleotides, which comprise known microsatellites (MS), located correctly by a tool. Known MS were found by RepeatMasker with a maximum mutation rate of 25%. False Positive Rate (FPR) is the number of nucleotide found in 1,000,000 base pairs of a random genome with the same length and nucleotide composition as the input genome; it is measured in base pair/mega base pair (bp/mbp). Precision is the ratio of the correctly located nucleotides to the total of this number and the number of nucleotides found in the random genome. F-measure is the harmonic mean of sensitivity and precision. Novel MS Content is the length (in mbp) of novel MS located by a tool. Percentage Coverage is the percentage of a genome made of MS. Time is measured in seconds (s) and memory in Megabytes (MB).

| Tool | F-measure (%) | Precision (%) | Sensitivity (%) | FPR (bp/mbp) | Novel MS Content (mbp) | Percent Coverage (%) | Time (s) | Memory (MB) |
| --- | --- | --- | --- | --- | --- | --- | --- | --- |
| *Arabidopsis thaliana* |  |  |  |  |  |  |  |  |
| Look4TRs | 71.3 | 97.7 | 56.1 | 92 | 3.4 | 3.3 | 55 | 2242 |
| MISA | 45.4 | 92.8 | 30.1 | 165 | 0.4 | 0.5 | 97 | 163 |
| MsDetector | 63.4 | 94.8 | 47.6 | 188 | 1.2 | 1.3 | 41 | 347 |
| Tantan | 51.0 | 38.7 | 75.0 | 8457 | 8.2 | 6.6 | 62 | 167 |
| TRF | 36.5 | 24.7 | 69.9 | 15197 | 5.5 | 3.8 | 199 | 203 |
| *Drosophila melanogaster* |  |  |  |  |  |  |  |  |
| Look4TRs | 83.3 | 97.4 | 72.7 | 350 | 6.4 | 5.8 | 103 | 2507 |
| MISA | 32.5 | 97.9 | 19.5 | 77 | 0.5 | 0.7 | 168 | 169 |
| MsDetector | 68.7 | 99.0 | 52.6 | 97 | 2.4 | 2.6 | 48 | 366 |
| Tantan | 77.1 | 76.4 | 77.9 | 4406 | 9.0 | 7.7 | 72 | 174 |
| TRF | 62.3 | 54.5 | 72.6 | 11098 | 7.5 | 6.5 | 357 | 218 |
| *Homo sapiens* |  |  |  |  |  |  |  |  |
| Look4TRs | 91.3 | 95.1 | 87.8 | 376 | 111.3 | 4.3 | 1003 | 3570 |
| MISA | 52.5 | 97.3 | 36.0 | 83 | 16.4 | 0.8 | 3863 | 1301 |
| MsDetector | 83.8 | 98.8 | 72.8 | 72 | 44.8 | 2.0 | 1020 | 2645 |
| Tantan | 71.2 | 60.3 | 87.1 | 4745 | 160.1 | 5.8 | 1609 | 1340 |
| TRF | 51.9 | 37.5 | 84.5 | 11647 | 128.9 | 4.8 | 7670 | 3934 |
| *Hordeum vulgare* |  |  |  |  |  |  |  |  |
| Look4TRs | 85.2 | 87.3 | 83.2 | 396 | 202.8 | 4.4 | 1733 | 6058 |
| MISA | 36.6 | 94.2 | 22.7 | 46 | 6.1 | 0.2 | 4071 | 3804 |
| MsDetector | 73.5 | 98.1 | 58.7 | 37 | 49.5 | 1.2 | 1506 | 7909 |
| Tantan | 55.8 | 43.8 | 77.0 | 3219 | 250.7 | 5.4 | 2525 | 4127 |
| TRF | 30.6 | 19.4 | 72.7 | 9880 | 148.3 | 3.3 | 6287 | 1205 |
| *Oryza sativa Japonica* |  |  |  |  |  |  |  |  |
| Look4TRs | 82.7 | 97.7 | 71.7 | 113 | 24.4 | 7.0 | 213 | 3117 |
| MISA | 51.6 | 97.6 | 35.1 | 56 | 1.2 | 0.5 | 305 | 228 |
| MsDetector | 71.7 | 98.8 | 56.3 | 44 | 6.1 | 1.2 | 127 | 490 |
| Tantan | 67.0 | 58.6 | 78.2 | 3665 | 33.5 | 5.4 | 194 | 236 |
| TRF | 45.4 | 32.5 | 75.5 | 10404 | 19.5 | 4.3 | 712 | 255 |
| *Plasmodium falciparum* |  |  |  |  |  |  |  |  |
| Look4TRs | 93.0 | 99.5 | 87.3 | 492 | 4.0 | 26.6 | 49 | 2435 |
| MISA | 65.0 | 97.5 | 48.7 | 1429 | 0.6 | 8.0 | 24 | 31 |
| MsDetector | 84.1 | 94.5 | 75.7 | 4949 | 2.2 | 17.6 | 7 | 41 |
| Tantan | 81.5 | 98.0 | 69.8 | 1624 | 2.0 | 16.1 | 12 | 20 |
| TRF | 70.9 | 60.2 | 86.1 | 63705 | 3.5 | 24.3 | 613 | 62 |
| *Sorghum bicolor* |  |  |  |  |  |  |  |  |
| Look4TRs | 79.6 | 96.3 | 67.8 | 123 | 23.6 | 3.7 | 219 | 2822 |
| MISA | 52.2 | 96.7 | 35.7 | 57 | 1.0 | 0.3 | 558 | 395 |
| MsDetector | 77.0 | 98.5 | 63.2 | 45 | 5.8 | 1.1 | 236 | 902 |
| Tantan | 61.7 | 50.4 | 79.4 | 3659 | 36.4 | 5.7 | 376 | 419 |
| TRF | 39.6 | 26.4 | 79.6 | 10404 | 25.9 | 4.1 | 1035 | 232 |
| *Zea mays* |  |  |  |  |  |  |  |  |
| Look4TRs | 84.3 | 90.7 | 78.8 | 168 | 92.0 | 4.5 | 767 | 4164 |
| MISA | 46.3 | 94.5 | 30.7 | 38 | 1.8 | 0.1 | 1735 | 1529 |
| MsDetector | 74.3 | 97.5 | 60.0 | 32 | 25.3 | 1.3 | 728 | 3414 |
| Tantan | 48.9 | 35.7 | 77.5 | 2911 | 136.4 | 6.6 | 1091 | 1651 |
| TRF | 24.2 | 14.4 | 76.2 | 9469 | 76.0 | 3.8 | 2871 | 580 |

### Contributions of this study

Our efforts have led to the following contributions:

- The Look4TRs software: This tool is the first application of using self-supervised hidden Markov models to detecting MS. Further, this is the first adaptive, auto-calibrating tool for discovering MS. Furthermore, Look4TRs, similar to MsDetector, is parameter-free, leading to consistent results across different studies. The C++ source code is available on GitHub (https://github.com/TulsaBioinformaticsToolsmith/Look4TRs) and as Supplementary Data Set 1.
- We applied Look4TRs to locating MS in the genomes of the following eight species: *Homo sapiens, Arabidopsis thaliana, Hordeum vulgare, Oryza sativa Japonica, Sorghum bicolor, Zea mays, Drosophila melanogaster, and Plasmodium falciparum*. Microsatellites located in the eight genomes by Look4TRs are available as the Supplementary Data Set 2–9.

### Evaluations on eight genomes

We evaluated Look4TRs and four related tools — MISA, MsDetector, Tantan, and TRF — on the eight genomes. Table 1 shows the results of these evaluations. Our goal is to design MS-detection tool with high sensitivity and low false positive rate. Recall that the F-measure combines the sensitivity and the precision measures. Therefore, the F-measure is the most important evaluation criterion. Look4TRs achieved the highest F-measure on all genomes with a clear margin of improvement over the second best performing tool, which was MsDetector on seven genomes and Tantan on one genome. The improvement ranged from 3.4% up to 15.9%, averaging 11.0% on the eight genomes. The improvement over TRF, which is one of the most widely-used tools for MS detection, averaged 106.4%. Similarly, the improvment over MISA, another widely-used tool, averaged 82.2%. Look4TRs outperformed Tantan by 33.5%, on average, on the eight genomes. Tantan is a recently developed tool based on hidden Markov models. These results indicate that Look4TRs achieves high sensitivity without compromising the false positive rate.

With respect to precision, Look4TRs was the most precise on two genomes — *Arabidopsis thaliana* and *Plasmodium falciparum*. The *Plasmodium falciparum* genome has very high AT percentage, representing a challenge to repeat-detection tools in general. Achieving the highest precision on this genome indicates that Look4TRs is highly adaptive to the input genome and is likely to detect MS successfully in other genomes with skewed nucleotide distributions. On the other six genomes, MsDetector was the most precise tool. MsDetector was more precise than Look4TRs by 1.1%–12.4%, averaging 4.8% on the six genomes.

Regarding sensitivity, Look4TRs was more sensitive or comparable to the most sensitive tool on four genomes; it came second on one genome and third on the other three genomes. On the four genomes, on which Look4TRs was not the most sensitive tool, the best performing tool (Tantan or TRF) was more sensitive than Look4TRs by 7.2%–33.7%, averaging 16.8% on these four genomes. Tantan was the most sensitive tool or comparable to most sensitive tool on seven genomes. The sensitivity of TRF was comparable to the sensitivities of the best performing tools on three genomes. TRF sensitivity came second on four genomes and third on the last genome. MsDetector sensitivities came fourth ranging from 47.6% to 75.5%. Finally, MISA was consistently the least sensitive tool (19.5%–48.7%). These results show that Look4TRs has reasonable sensitivity.

Look4TRs had the third lowest FPRs on six genomes and the lowest FPRs on two genomes. Even though it came third on six genomes, the FPRs of Look4TRs were very reasonable (92–492 bp/mbp). MISA and MsDetector has the lowest FPRs in general, whereas TRF had the highest.

With respect to the novel MS content, Look4TRs was first on one genome, second on three genomes, and third on the other four genomes. Given, the reasonable sensitivity and the low FPR of Look4TRs, these potential novel MS are likely true tandem repeats. Table 2 displays example novel MS found by Look4TRs.

One application of MS-discovery tools is the estimation of MS size or MS content in a genome. On the five plants genomes, the MS contents were between 3.3% (*Arabidopsis thaliana*) and 7.0% (*Oryza sativa Japonica*). MS found by Look4TRs in the *Homo sapiens* genome comprise 4.3% and those found in the *Drosophila melanogaster* genome comprise 5.8%. Let’s consider the *Plasmodium falciparum* genome, on which Look4TRs estimated the MS content to be 26.6%. Recall that the AT-content of this genome is 80%. Because Look4TRs has the lowest FPR, which was calculated on a synthetic genome with 80% AT-content, these predicted satellites are unlikely to be false positives. This genome has another interesting feature; it does not include transportable elements^25^. Is it possible that the abundance of tandem repeats compensate for the absence of the interspersed ones in the *Plasmodium falciparum* genome? This question remains to be answered!

With respect to time, Look4TRs was among the fasted tools; it took 0.8–28.9 minutes to process one of the tested genomes. Look4TRs time requirements were measured using eight threads. By default, Look4TRs is multi-threaded to take advantage of the concurrency built in personal computers. Recall that we ran the other tools with their default parameters, just as an average user would use them. It is possible that their time requirements are less than what we reported if they have parameters controlling concurrency or ran using the GNU “parallel” utility. However, using the “parallel” utility will increase the memory requirements, which we discuss next. The memory requirements of Look4TRs were among the highest. It used 1–6 GB of memory. However, the required memory is readily available on any recent personal computer.

**Table 2.**
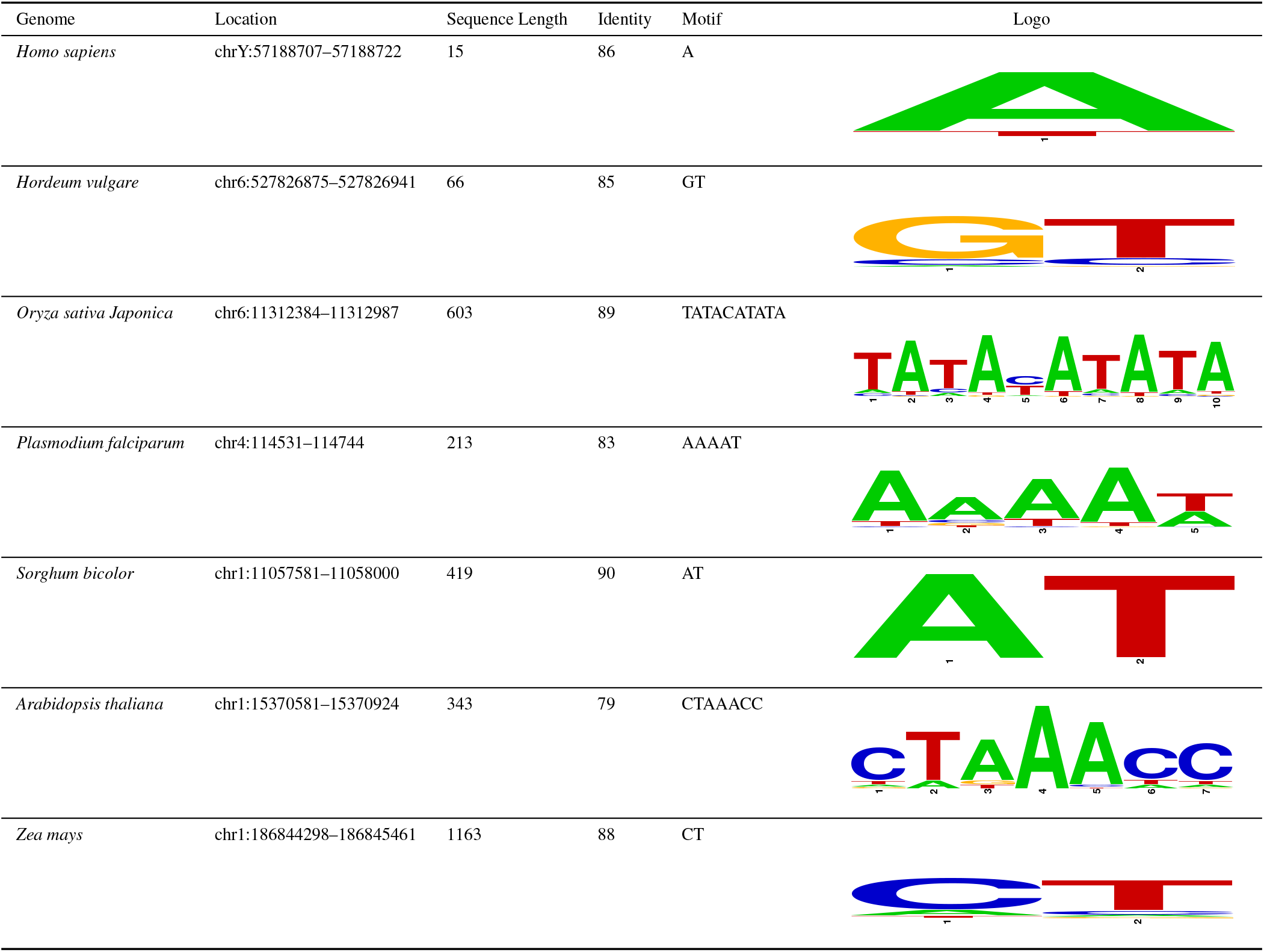
Example novel microsatellites detected by Look4TRs. Sequence Length is the length of a predicted region. Identity represents the alignment identity score between the located region and the theoretical perfect MS consisting of multiple exact copies of the motif. Motif represents the repeated motif found by Look4TRs in a region. Logos of these motifs are shown under the column labeled “Logo” and were produced by WebLogo^26^.

| Genome | Location | Sequence Length | Identity | Motif | Logo |
| --- | --- | --- | --- | --- | --- |
| *Homo sapiens* | chrY:57188707–57188722 | 15 | 86 | A |  |
| *Hordeum vulgare* | chr6:527826875–527826941 | 66 | 85 | GT |  |
| *Oryza sativa Japonica* | chr6:11312384–11312987 | 603 | 89 | TATACATATA |  |
| *Plasmodium falciparum* | chr4:114531–114744 | 213 | 83 | AAAAT |  |
| *Sorghum bicolor* | chr1:11057581–11058000 | 419 | 90 | AT |  |
| *Arabidopsis thaliana* | chr1:15370581–15370924 | 343 | 79 | CTAAACC |  |
| *Zea mays* | chr1:186844298–186845461 | 1163 | 88 | CT |  |

In sum, Look4TRs is able to maintain a high sensitivity that does not interfere with its FPR. Tantan and TRF achieve higher sensitivities than Look4TRs on three genomes — *Arabidopsis thaliana, Oryza sativa Japonica*, and *Sorghum bicolor*. However, Tantan and TRF have FPRs that are consistently an order of magnitude greater than the FPR obtained by Look4TRs. This indicates that Tantan and TRF simply predict more sequences to be tandem repeats, which leads to many false positives that are not produced by Look4TRs. This also explains why Tantan and TRF achieve higher novel MS contents. Simply, Look4TRs is able to find many simple repetitive sequences without compromising the FPR. On the other end of the spectrum, MISA and MsDetector had the lowest FPRs; however, these low rates resulted in lower sensitivities. The FPR of Look4TRs is slightly higher than those achieved by MISA and MsDetector, but Look4TRs sensitivity is much higher. Therefore, Look4TRs is the best available tool that provides high sensitivity while maintaining low FPR. This conclusion is supported by the consistent high F-measure values achieved by Look4TRs on the eight tested genomes. These results suggest that Look4TRs can annotate MS in the newly sequenced genome accurately.

## Methods

### Overview

Here, we describe our software tool, Look4TRs — the source code is available on GitHub (https://github.com/TulsaBioinformaticsToolsmith/Look4TRs) and as Supplementary Data Set 1. It is designed to search an entire genome for microsatellites (MS). The software tool consists of the following four modules:

- The scoring module: It converts a sequence of nucleotides to a sequence of scores. A score indicates whether or not the corresponding nucleotide is in a repetitive region.
- The training module: The module is a self-supervised system which generates a random synthetic chromosome that has the same nucleotide composition as the largest chromosome of the input genome. This random chromosome has synthetic MS inserted into known locations. For every choice of parameters that affect the scoring module, a hidden Markov model (HMM) is trained on the random chromosome and the HMM’s output is compared to the true locations of the synthetic repeats. The parameters which produces the best output will be selected to train the HMM that will scan the entire genome.
- The scanning module: Using the trained HMM, the Viterbi algorithm delineates the final repetitive regions.
- The motif-discovery module: The module attempts to identity the repeated motif in a region. To start, candidate motifs are determined. After that, a sequence is generated from each motif by appending multiple copies of it in tandem, resulting in a sequence that has the same length as the predicted region. This sequence is then compared to the predicted region. The motif of the most similar sequence to the predicted region is selected.

Next, we illustrate how each module works.

### The scoring module

This module is designed to generate scores, which distinguish nucleotides found in MS from those found outside repetitive regions. It is an *adaptive* component, accounting for the nucleotide bias found in a species such as the *Plasmodium falciparum* — %80 AT-content. The idea of utilizing sequence composition information was inspired by Achaz, et al.^20^ We have applied a similar idea successfully in Red, which is a tool for finding tandem and interspersed repeats^21^. In Look4TRs, a nucleotide score reflects how repetitive a k-mer starting at this nucleotide is in the small surrounding region/window. The observed and the expected counts of a k-mer are required to calculate the score (Equation 5).

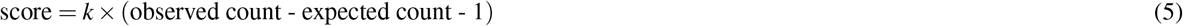

If the score is negative, it is set to zero. To calculate the observed count, the k-mer is counted in the window. The expected count of the same k-mer is estimated using a Markov chain of order zero trained on the sequence within the window. The final score of a nucleotide is the highest score resulting from trying different values of k, e.g. 4–10. To process long sequences efficiently, a k-mer is represented as a quaternary number. Horner’s rule is used for calculating the quaternary numbers of a long sequence efficiently^21^. The counts of k-mers in the window are stored in a hash table — a separate table for each value of k. Because the module processes consecutive nucleotides, there is no need to recount all k-mers in the window centered on the next nucleotide. Instead, the count of the k-mer starting at the first nucleotide of the current window is decremented by one and the count of the k-mer starting at the last nucleotide of the next window is incremented by one.

### The training module

The core of this module is a self-supervised HMM. HMMs are instances of supervised learning, requiring the availability of labels. The labels in the case at hand are the locations of known MS. A self-supervised algorithm can generate its own labels. Earlier, we successfully invented self-supervised systems for predicting enhancers^27^, masking repeats^21^, clustering DNA sequences^28^, and predicting the identity score of two sequences in linear time^29^. Next, we discuss (i) how the labels are generated, (ii) how the HMM is trained, and (iii) how the module auto-calibrates itself.

#### Generating the labels

Using a Markov chain of order zero trained on the largest chromosome of the input genome. A random synthetic chromosome is generated so that it has the same nucleotide composition as the real chromosome. One reason for choosing the largest chromosome is that the largest chromosome will typically have the most data needed for training. Another reason for using the largest chromosome is that we would like to minimize any variability that may arise from training on different chromosomes. All what is needed is the percentage of each of the four nucleotides in the input genome. Percentages found in the largest chromosome should be very similar to those found in the entire genome. We made sure that the same synthetic chromosome is generated from the same real chromosome every time the program is executed, i.e. multiple runs of Look4TRs on the same genome will produce the same results. Next, MS are generated, mutated, and inserted into the synthetic chromosome, while their locations are recorded. These tandem repeats are based off of sequences found in the original chromosome and make up 5% of the random chromosome. To generate a tandem repeat, a short motif is selected randomly from the real chromosome; then it is repeated multiple times; the synthetic microsatellite is mutated according to a mutation rate between 0% and 25% distributed uniformly. Algorithm 1 provides the pseudo code of how the labels required for training the HMM are generated.

##### Algorithm 1

Generating semi-synthetic microsatellites (MS), which serve as the labels needed to train the hidden Markov model, making Look4TRs a self-supervised system.

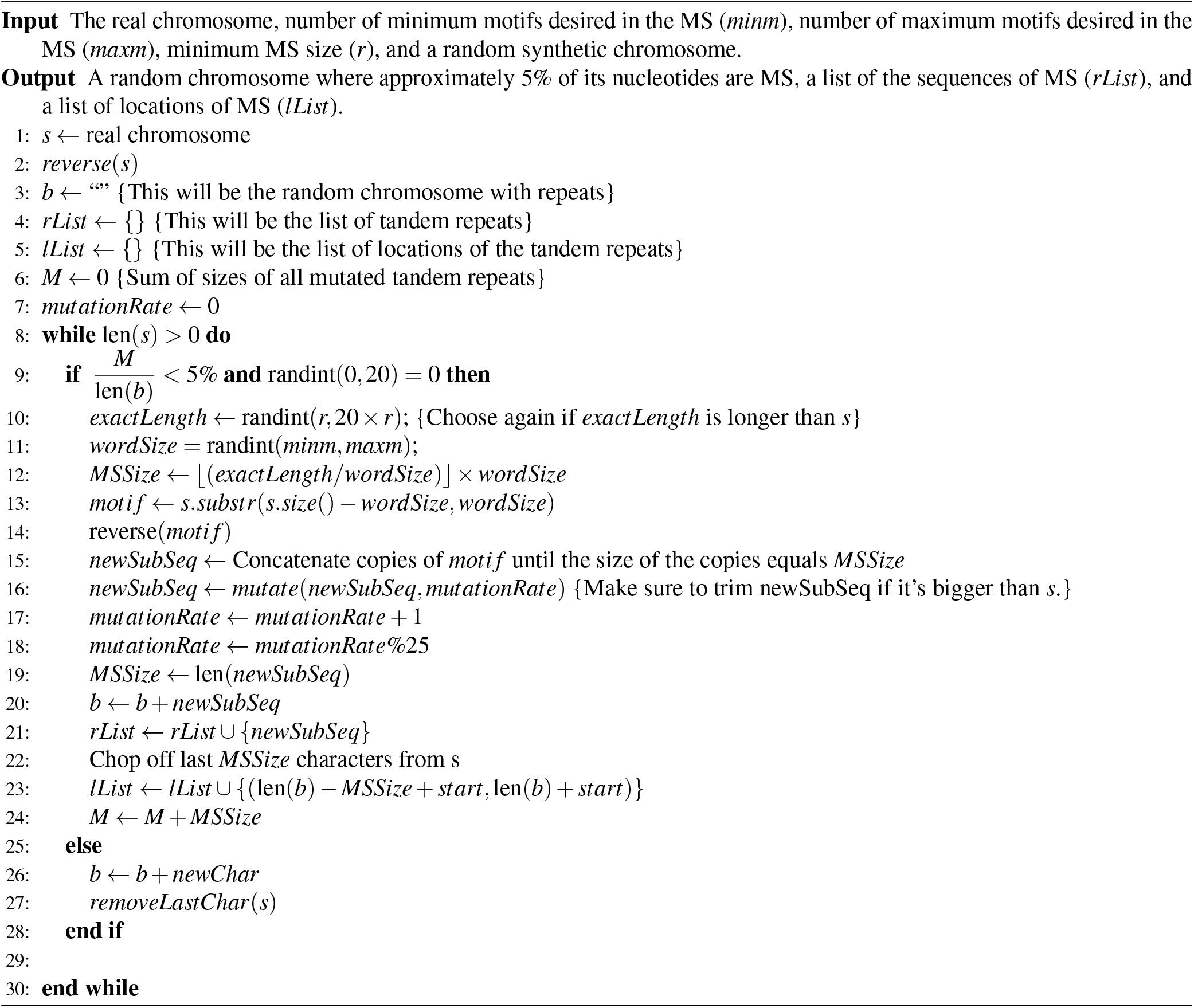

#### Training the HMM

The HMM is trained on the scores of the generated MS and the scores of the regions in between. Earlier, we implemented an HMM for detecting all types — tandem, interspersed, and low-complexity regions — of repeats^21^. Here, we utilized the same model in detecting MS. An HMM consists of a set of states and three types of probabilities known as the prior, the transition, and the output probabilities. Our HMM has 20 states: 10 positive states representing MS and 10 negative states representing non-repetitive regions. The number of states has a minimal effect on the performance as long as it is large enough. A positive state represents a range of scores, which have the same logarithmic value, found in a repetitive region. Similarly, a negative state represents a range of scores found outside the MS. Thus, each score has two potential states associated with it. In our implementation, the scores were preprocessed by taking their logarithms (base 2). At this point, the log score indicates the number labeling its state; the index of the score indicates the sign; for example, a log score of 2 in a repetitive region is considered to be generated by the +2 state. The HMM is trained on the MS and the non-MS regions by counting the number of transitions between every two consecutive states, as well as the number of each state that begins an MS region or a non-MS region. Next, these counts are normalized into probabilities. The output probabilities are set to 1 for all states because a state outputs one log score only.

#### Auto-calibrating the training module

After generating a synthetic chromosome, multiple scoring modules are used for generating scores for training multiple HMMs. The scoring module takes the following parameters: (i) a minimum k-mer size, (ii) a maximum k-mer size, and (iii) the size of the half-window. Given a lower and upper bounds for the k-mer size, a list of pairs of valid minimum and maximum k-mer sizes is generated. As for the half-window size, four multiples of the initial region size (20 base pairs) are tested 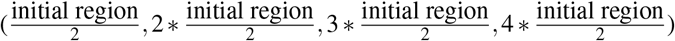. Every combination of minimum k-mer size, maximum k-mer size, and half-window sizes is used for training an HMM. The HMM — using the Viterbi algorithm — outputs a list of tandem repeat locations it detects in the synthetic chromosome. These HMM locations can be compared to the actual locations to obtain the F-measure, which combines the sensitivities and the precision measures (Equations 4, 1, and 2). A high F-measure indicates high sensitivity and low false positive rate. The parameters which lead to the HMM with the highest F-measure will be chosen to train the final HMM in the scanning module. The benefit is that this reduces the responsibility of the user to pick optimal parameters since *this module auto-calibrates and chooses the best parameters itself*.

### The detection module

Once the auto-calibration process in complete and the parameters have been determined, the final HMM is trained. The Viterbi algorithm, which is an instance of dynamic programming, is used for determining the most probable series of states that generated a sequence of log scores. A sequence of positive states represents a repetitive region, and a sequence of negative states represents a non-repetitive region.

### The motif-discovery module

This module aims at identifying the repeated motif in a tandem repeat utilizing a recently developed algorithm for predicting the similarity between two sequences efficiently.

#### Predicting identity scores using k-mer features

Traditionally, sequence identity is calculated using a quadratic-time alignment algorithm. Aligning two sequences can take a long time, especially on long sequences. Alternatively, k-mers can be counted; then different features are extracted from these counts. First, we surveyed and evaluated more than 30 k-mer features^30^, which can be calculated in linear time. Few of these features — and their squared versions and multiplicative combinations of every two features — are combined in a regression model for predicting the sequence identity scores. Look4TRs’s regression model is based on a General Linear Model (GLM).

A GLM is an instance of supervised learning; it learns the optimal weights associated with the input features in order to predict the labels. If the labels are classes, then the task is classification. If the labels represent a continuous variable, then the task is regression; this is the case here. GLMs have been applied broadly in bioinformatics. Previously, we have applied GLMs to ranking the quality of predicted protein structures^31–33^, filtering out spurious MS^22^, and predicting the similarity between two DNA sequences^28, 29^. We have devised a similar GLM-based classifier in MeShClust^28^, which is a tool for clustering DNA sequences, and a similar GLM-based regression model in FASTCAR^29^, which is a tool for approximating the identity score between two DNA sequences in linear time.

#### Generating training data for the GLM

The utilized GLM is an instance of self-supervised learning because the labels are generated automatically. The GLM is trained on 1000 sequence pairs (the training set), and tested on different 1000 sequence pairs (the testing set). The semi-synthetic MS generated for training the HMM are also used in training the GLM. First, we find all words of size 1–10 nucleotides that occur in one MS region. For each word, a synthetic sequence is constructed by appending the word in tandem, i.e. a synthetic tandem repeat. The synthetic sequence has the same length as the original MS. The MS and a synthetic sequence makes one pair. Note that one MS region can be paired with several synthetic sequences, each of which is due to one word found in the original MS. The identity score — the label — of each pair of sequences is calculated by the alignment algorithm.

#### Selecting features

Once the 2000 sequence pairs are accumulated, we apply a feature-selection algorithm to guard against over-fitting, in which a model tends to memorize the training examples rather than to learn a general concept. The feature selection algorithm uses a greedy approach that selects the best feature at each step. This feature-selection strategy is the same approach we utilized in FASTCAR^29^, whereas MeShClust’s classifier has four predetermined features and does not utilize this feature selection algorithm. Doing these steps accumulates features that improve the mean error (the absolute difference between the predicted identity score and that due to the alignment algorithm) at every step on the testing set, which is different from the training set.

##### Algorithm 2

Discovering a motif in microsatellites

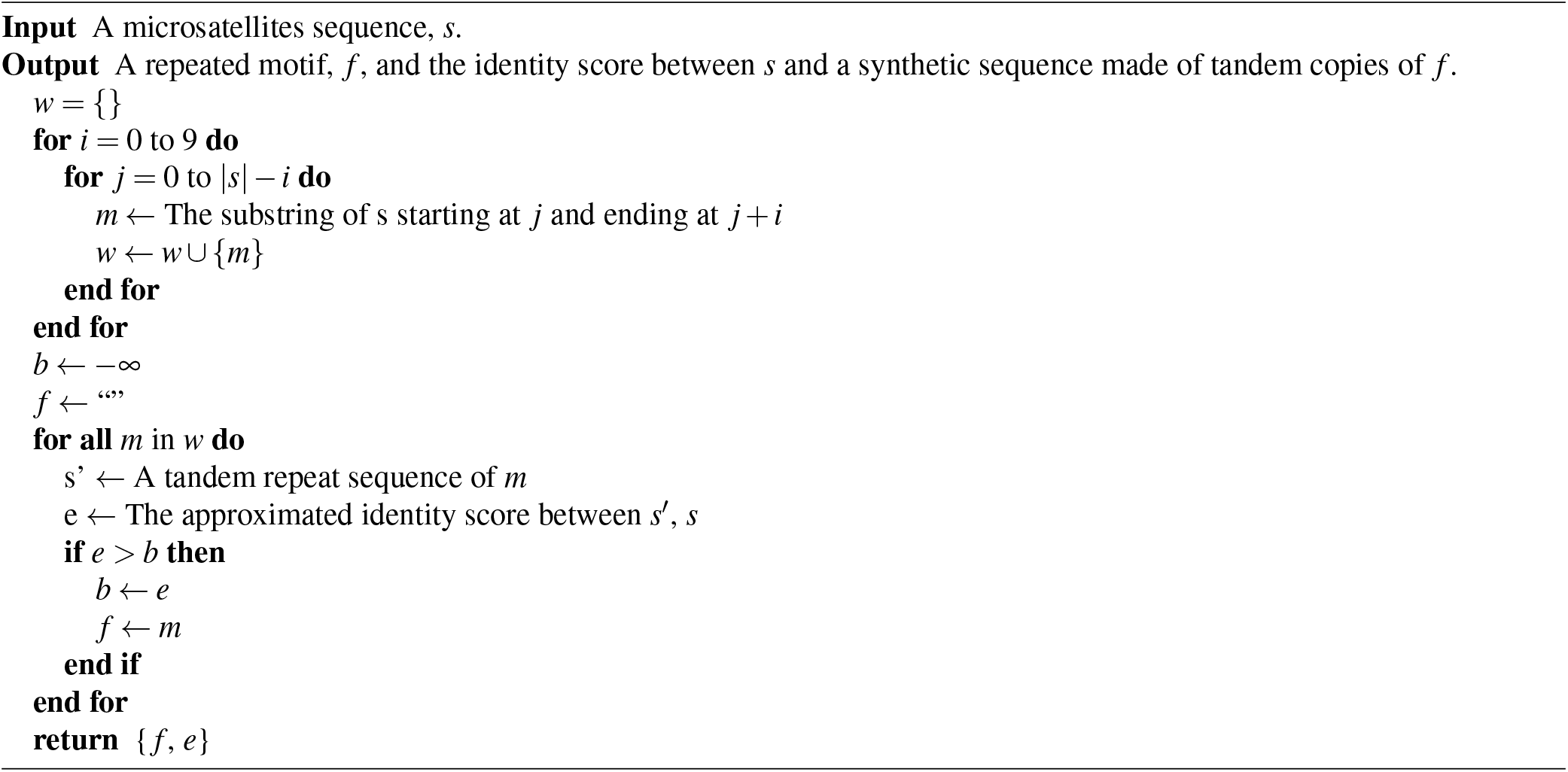

After the informative features are selected, the GLM is trained and tested. Next, we discuss how the regression model is utilized in identifying the repeated motif in potential MS.

#### Finding repeated motifs in microsatellites

In Algorithm 2, we outline the algorithm for finding a motif in microsatellites. As we have done earlier for training the GLM, we find all words of size 1-10 nucleotides that occur in the input sequence. For each word, a synthetic sequence is constructed by appending the word in tandem. Next, the identity score between the original and the synthetic sequences is calculated by the regression model. After that, we select the word resulting in the highest identity; if there is a tie, the shortest word is picked.

### Related tools

To run TRF, we reviewed the documentation provided, and attempted to use the most reasonable parameters for microsatellites. We used 2, 7, 7 as the match, the mismatch, and the indel parameters, as recommended. For the match probability and the indel probability, we again used the recommended parameters of 80 and 10. The minimum score parameter was set to be 50, as recommended in the author’s paper^17^. For the “maxperiod” parameter, we used 10. Lastly, we used the “-l6” flag to limit the search size to 6 mbp instead of the entire chromosome. We ran Tantan using its default parameters, except on the *Plasmodium flaciparum* genome (80% AT-content) where the increased AT matrix was utilized. MsDetector was applied without parameters on all genomes except the *Plasmodium flaciparum* genome, on which we ran a version optimized for this genome. MISA was run using the default parameter file that is included in the source code and on the server. The unit_size-min_repeats pairs used are 1-10, 2-6, 3-5,4-5, 5-5, and 6-5. The max difference between 2 tandem repeats is set to be 100.

### Data

Repeats that diverged by 25% at most from the consensus sequence comprise the ground truth, which was obtained by RepeatMasker (http://www.repeatmasker.org) using Repbase^23^,^24^. We evaluated the above mentioned tools on the sequences of the following eight genomes:

- *Homo sapiens* (Hg38): http://hgdownload.cse.ucsc.edu/goldenPath/hg38/bigZips/
- *Arabidopsis thaliana* (TAIR10): http://plants.ensembl.org/Arabidopsis_thaliana/Info/Index
- *Hordeum vulgare* (HvIbscPgsbV2): http://plants.ensembl.org/Hordeum_vulgare/Info/Index
- *Oryza sativa Japonica* (IRGSP1): http://plants.ensembl.org/Oryza_sativa/Info/Index
- *Sorghum bicolor* (SorghumBicolorV2): http://plants.ensembl.org/Sorghum_bicolor/Info/Index
- *Zea mays* (ZeaMaysAGPv4): http://ensembl.gramene.org/Zea_mays/Info/Index
- *Drosophila melanogaster* (Dm6): http://hgdownload.soe.ucsc.edu/downloads.html#fruitfly
- **Plasmodium falciparum** (Pf3d7): http://www.sanger.ac.uk/resources/downloads/protozoa

## Conclusion

Multiple large-scale sequencing projects are underway. These projects will result in the sequences of hundreds of thousands of species. An important component of the majority of these genomes is simple tandem repeats — microsatellites (MS) in particular. The advances in the development of software for detecting MS did not keep up with the rapid advancements in sequencing technology. Based on performance, the currently available tools fall in two categories. The first category includes tools that are very sensitive; however, they have high False Positive Rates (FPRs). The second category includes tools that have low FPRs, but not very sensitive. None of these tools, in their default mode, takes into account the characteristics of the input genome. In our opinion, this lack of adaptability is the main limiting factor to the performances of the currently available tools. Adjusting the parameters manually on each new genome is impractical because there will be thousands, even hundreds of thousands, of new genomes.

Therefore, there is an immediate need for a new, adaptive tool that balances sensitivity and FPR. To this end, we propose Look4TRs, which utilizes self-supervised hidden Markov models for the first time in discovering MS. Additionally, Look4TRs has a scoring module that considers the nucleotide composition of the genome of interest. Further, Look4TRs auto-calibrates itself, relieving the user from adjusting its parameters and ensuring consistent results across different studies on the same genome.

We evaluated Look4TRs on eight genomes using different evaluation criteria. The results show that Look4TRs represents improvements over the currently available tools. Based on the F-measure, which takes into account both of the sensitivity and the FPR, Look4TRs outperforms TRF and MISA — the most widely-used tools for discovering MS — by very wide margins. Look4TRs also outperforms the recent related tools with narrower margins, however. We applied Look4TRs to estimating the percentages of MS in eight genomes. Interestingly, MS comprise 7.0% of the rice genome; this percentage is the highest among the five plant genomes we analyzed. In the human genome, MS comprise 4.3%. In sum, Look4TRs represents methodological advancement in the field of MS discovery, leading to a novel, adaptive tool that balances sensitivity and FPR.

## Acknowledgments

This research was supported mainly by funds from the Oklahoma Center for the Advancement of Science and Technology [PS17-015] and in part by internal funds provided by the College of Engineering and Natural Sciences and the Tulsa Undergraduate Research Challenge (TURC) Program at the University of Tulsa. We would like to thank Joseph Valencia for coding the global alignment algorithm and Robert Geraghty for coding the GLM.

## Author contributions statement

H.Z.G contributed the following: (i) conceived the idea, (ii) designed the software, (iii) implemented the scoring module, (iv) implemented the hidden Markov model, (v) designed the experiments, (vi) analyzed the results, and (vii) wrote the manuscript. A.V. implemented the auto-calibration feature, the generation of the synthetic chromosome, and the motif-discovery module; he conducted the experiments, produced the results, and contributed to the writing of the Methods and the Results Sections. B.T.J implemented the sequence identity approximation program and wrote the corresponding subsection of the Methods Section. V.D.W implemented an early prototype of the system. All authors read the manuscript.

## Data availability

The source code of Look4TRs is available on GitHub (https://github.com/TulsaBioinformaticsToolsmith/Look4TRs) and as Supplementary Data Set 1. The microsatellites found by Look4TRs in the eight genomes are available as Supplementary Data Set 2–9.

## Supplementary Information

**Supplementary Data Set 1 — C++ source code of Look4TRs**

A compressed file (.tar.gz) containing the C++ source code of Look4TRs as well as the manual and instructions on how to compile and run the program.

**Supplementary Data Set 2 — Microsatellites of Arabidopsis thaliana**

A compressed file (.tar.gz) containing the microsatellites located by Look4TRs in the genome of Arabidopsis thaliana.

**Supplementary Data Set 3 — Microsatellites of Drosophila melanogaster**

A compressed file (.tar.gz) containing the micro satellites located by Look4TRs in the genome of Drosophila melanogaster.

**Supplementary Data Set 4 — Microsatellites of Homo sapiens**

A compressed file (.tar.gz) containing the microsatellites located by Look4TRs in the genome of Homo sapiens.

**Supplementary Data Set 5 — Microsatellites of Hordeum vulgare**

A compressed file (.tar.gz) containing the microsatellites located by Look4TRs in the genome of Hordeum vulgare.

**Supplementary Data Set 6 — Microsatellites of Oryza sativa Japonica**

A compressed file (.tar.gz) containing the microsatellites located by Look4TRs in the genome of Oryza sativa Japonica.

**Supplementary Data Set 7 — Microsatellites of *Plasmodium falciparum***

A compressed file (.tar.gz) containing the microsatellites located by Look4TRs in the genome of *Plasmodium falciparum*.

**Supplementary Data Set 8 — Microsatellites of Sorghum bicolor**

A compressed file (.tar.gz) containing the microsatellites located by Look4TRs in the genome of Sorghum bicolor.

**Supplementary Data Set 9 — Microsatellites of Zea mays**

A compressed file (.tar.gz) containing the microsatellites located by Look4TRs in the genome of Zea mays.

